# The effect of speech masking on the human subcortical response to continuous speech

**DOI:** 10.1101/2024.12.10.627771

**Authors:** Melissa J. Polonenko, Ross K. Maddox

## Abstract

Auditory masking—the interference of the encoding and processing of an acoustic stimulus imposed by one or more competing stimuli—is nearly omnipresent in daily life, and presents a critical barrier to many listeners, including people with hearing loss, users of hearing aids and cochlear implants, and people with auditory processing disorders. The perceptual aspects of masking have been actively studied for several decades, and particular emphasis has been placed on masking of speech by other speech sounds. The neural effects of such masking, especially at the subcortical level, have been much less studied, in large part due to the technical limitations of making such measurements. Recent work has allowed estimation of the auditory brainstem response (ABR), whose characteristic waves are linked to specific subcortical areas, to naturalistic speech. In this study, we used those techniques to measure the encoding of speech stimuli that were masked by one or more simultaneous other speech stimuli. We presented listeners with simultaneous speech from one, two, three, or five simultaneous talkers, corresponding to a range of signal-to-noise ratios (SNR; Clean, 0, −3, and −6 dB), and derived the ABR to each talker in the mixture. Each talker in a mixture was treated in turn as a target sound masked by other talkers, making the response quicker to acquire. We found consistently across listeners that ABR wave V amplitudes decreased and latencies increased as the number of competing talkers increased.

**Significance statement:** Trying to listen to someone speak in a noisy setting is a common challenge for most people, due to auditory masking. Masking has been studied extensively at the behavioral level, and more recently in the cortex using EEG and other neurophysiological methods. Much less is known, however, about how masking affects speech encoding in the subcortical auditory system. Here we presented listeners with mixtures of simultaneous speech streams ranging from one to five talkers. We used recently developed tools for measuring subcortical speech encoding to determine how the encoding of each speech stream was impacted by the masker speech. We show that the subcortical response to masked speech becomes smaller and increasingly delayed as the masking becomes more severe.

## Introduction

Speech is foundational to communication for hearing people. While some speech is heard under quiet conditions like a living room or office, noisy listening conditions like busy streets, crowded restaurants, or transit are daily scenarios that make understanding speech difficult. Masking is the phenomenon by which this noise negatively impacts the processing of speech. Overcoming masking to understand speech is built on an expansive neurophysiological network that begins in the cochlea and involves numerous interconnected regions in the brainstem, thalamus, and cortex. While many studies have investigated masking, little is known about the neural encoding of masked, naturally uttered speech in human listeners, especially in the earlier subcortical stages of the auditory system. The present study was aimed at addressing that gap.

Masking of speech has been extensively studied through psychophysical means for the better part of a century (Licklider, 1948; Miller and Licklider, 1950). These behavioral studies have described the perceptual phenomena of masking in fine detail, including distinct (but typically co-occurring) types of masking, such as energetic and informational masking (Brungart et al., 2001). There are also well-known attributes of stimuli or listening scenarios that offer some release from masking, such as spatial separation (Freyman et al., 2001) or the addition of visual cues (Sumby and Pollack, 1954).

Masking’s effects are often worsened in people with hearing loss and those using hearing aids and cochlear implants (Working Group on Speech Understanding and Aging, 1988; Fu and Nogaki, 2005), making its study clinically important as well. In some specific instances, behavioral masking effects can be closely tied to the underlying physiology, such as tone-in-noise paradigms designed to assess cochlear tuning in humans (Oxenham and Shera, 2003). But generally speaking, it is difficult to infer masking’s specific neurophysiological effects from behavioral data (which by their nature depend on the entire auditory system), especially for natural speech stimuli.

Neural studies of masking are less common. In humans, cortical responses to masked natural speech have been studied to better understand selective auditory attention (Mesgarani and Chang, 2012; Ding and Simon, 2013; O’Sullivan et al., 2014). Studies aimed at subcortical effects of speech masking have used repeated single syllables in multi-talker babble noise (Song et al., 2011). There has also been animal work using conspecific stimuli to investigate aspects of masking, such as masker type (Narayan et al., 2007) and spatial separation (Maddox et al., 2012), but these are harder to link to human perception.

There is a clear need for linking our understanding of the neural and perceptual effects of masked speech. Hearing aids can restore audibility for people with hearing loss, but speech understanding, especially in noise, remains a challenge for many (Davidson et al., 2022). Identifying the underlying neural causes may help us understand some of the variability across listeners with similar hearing loss configurations, and may also aid in developing and fitting better signal processing algorithms. Even people with clinically normal hearing thresholds can have significant difficulty understanding speech under noisy conditions. Multiple hypotheses exist regarding the neural and cochlear causes of such listening challenges (Bharadwaj et al., 2014; Carney, 2018), but a physiological test that accurately predicts speech-in-noise perception in humans with normal audiograms has been elusive (Prendergast et al., 2015). Many of these attempts have comprised responses to transient stimuli (e.g., clicks, tone bursts, repeated syllables), sometimes in the presence of masking noise, which bear little resemblance to natural speech.

The goal of this paper was to better understand masking’s effects on the subcortical neural encoding of naturally uttered speech in human listeners. To do this we leveraged our recently developed method for determining the auditory brainstem response (ABR) to speech (Polonenko and Maddox, 2021a), which is built on the temporal response function (TRF) framework (Lalor and Foxe, 2010).

Whereas our previous work was aimed at encoding of single talkers, here we determined the ABR (as a TRF reflecting subcortical response components) to speech in quiet as well as in the presence of varying numbers of other talkers. We found robust trends in the latency and amplitude of the responses as the number of talkers (and thus level of masking) increased. We also assessed how quickly those masking effects could be ascertained in individual listeners, which is critical for any measure that is a potential candidate for clinical use.

## Methods

### Human participants

All experimental methods were approved by the University of Rochester Research Subjects Review Board. Participants gave informed consent before participating, were paid for their time, and had confirmed hearing thresholds of 20 dB HL or better at octave frequencies from 500 to 8000 Hz in both ears. Twenty-five participants were recruited with ages from 19–37 years (mean ± SD of 23.4 ± 5.5 years) with 16 identifying as female and 9 as male.

### Behavioral task

While peaky speech preserves natural speech’s spectrotemporal structure and subjectively sounds very similar, it was important to confirm that masking effects were also similar (see next section for details of peaky speech construction). We conducted a listening task with speech-on-speech masking so that we could compare the speech reception thresholds of natural and peaky speech.

We used the standard coordinate response measure (CRM) sentences as target speech (Brungart, 2001), in a task based on Gallun et al. (2013). Sentences were randomly chosen from any of the male talkers and had the callsign “Charlie.” The targets were presented at 65 dB SPL over Etymotic ER-2 earphones. The masker sounds were speech randomly selected from 150 ten-second segments from the five stories used as stimuli in the EEG task (described in the next section). They were then shortened to be the same length as the CRM sentence. Each masker was a combination of segments from three distinct talkers from the set of five added together. The sound level was limited to 80 dB SPL, which meant that the minimum (most difficult) target-to-masker ratio (TMR) tested was −14.5 dB.

Before testing, participants were trained to make sure they understood the task. This was accomplished by presenting target sentences with no masker. After each response, a binomial test was performed on the responses so far under the null assumption that the responses were random. Each participant moved on to the main task once that assumption could be rejected at *p* =.05 for both natural and peaky speech. All participants passed training easily, in 3–8 trials per stimulus type.

Speech reception thresholds for the natural and peaky speech targets were determined through an adaptive track in which the masker level was varied. There was a random wait of one to four seconds between trials. The track started with 5-up-1-down for three reversals, and then became a 1-up-1- down track for eight reversals. Those latter eight were averaged to estimate the 50% correct point, where a correct trial was defined as choosing the correct color and number combination. With four color choices and eight numbers, chance level was one in thirty-two, or 3.1%.

Within each of six blocks, an adaptive track was run for each stimulus type. The final threshold for each stimulus type was taken as the average of the thresholds from the six blocks. Trials from the tracks were interleaved randomly, but the tracks were updated separately. We also ensured that one tracker never got more than two trials ahead of the other. Four breaks of a minimum of 15 s were forced during the experiment, and participants were allowed to take other breaks *ad libidum*, including extending the forced breaks.

### EEG

#### Stimuli

Five audiobooks, read by five different narrators, were downloaded from the open source LibriVox Collection: *Alice’s Adventures in Wonderland* (Carroll, 2020), *Peter Pan* (Barrie, 2017), *The Adventures of Pinocchio* (Collodi, 2012), *The Time Machine* (Wells, 2011), and *The Wonderful Wizard of Oz* (Baum, 2007). All narrators were male, since previous work has shown that peaky speech narrated by folks with lower f0s yields larger responses (Polonenko and Maddox, 2021a, 2024). Each story was filtered so that its long-term power spectrum matched the average spectrum of all five stories (Shan et al., 2024). Responses were then converted to peaky speech in the same manner as prior work (Polonenko and Maddox, 2021a). Briefly, converting natural speech to peaky speech involves the following steps:

1. Identify the times of glottal pulses (present during vowels and voiced consonants) using Praat (Boersma and Weenink, 2024) controlled in Python with Parselmouth (Jadoul et al., 2018).
2. Create a dynamic cosine whose phase advances by 2π between each glottal pulse, such that its instantaneous frequency matches that of the speech’s fundamental frequency. The amplitude of the cosine is determined from the spectrogram of the original speech.
3. Repeat step 2 for all harmonics (integer multiples of the fundamental frequency), with the exception that the frequency of each cosine is 2π*k* for the *k*^th^ harmonic. Add the fundamental and all harmonics together.
4. Combine the synthesized speech (which contains only the voiced portions of the speech) with the unvoiced portions of the original speech.

We created the different SNR conditions by mixing talkers from different stories together, with each talker always at the same overall level as the others. By mixing two talkers together, each talker was masked by one other talker at the same level, so the SNR was 0 dB (signal and noise were the same). For three talkers, one talker was masked by two others (doubling the noise energy), so the SNR was −3 dB. An SNR of −6 dB was created from five talkers (each one masked by four others). In general, the SNR could be calculated as −10 log_10_(*N* − 1), where *N* was the number of talkers. Single talkers were also presented in isolation, with that condition labeled as “Clean.” Combinations of narrators were created so that each story was presented with equal frequency over the course of the experiment. Each talker was presented diotically at 65 dB SPL, meaning that with five talkers the level was 72 dB SPL. This was deemed in piloting to be the highest level that participants considered comfortable. Trials from each SNR condition were recorded for 30 minutes, meaning there were 6, 12, 18, and 30 minutes of each story presented for the clean, 0, −3 and −6 dB SNR conditions.

An advantage of our speech presentation paradigm is that the response to all talkers in a mixture could be calculated separately and then averaged. By doing so, even though only 30 minutes of data were recorded from each SNR condition, there were 30, 60, 90, and 150 minutes of total speech presented for the clean, 0, −3 and −6 dB SNR conditions. This effective increase in recording time as SNR decreased led to reduced noise in the resulting waveforms, largely ameliorating the problem of smaller amplitude responses in those conditions (see Results).

Stimuli were presented in 10 s epochs with order randomized. Experiments were controlled with a python script built on the expyfun module (Larson et al., 2014). Stimuli were played using an RME Babyface Pro sound card at 48,000 samples / s, and presented over Etymotic Research ER-2 tube earphones.

### Recording

Participants were seated in reclining chair in a darkened sound-treated room. They were given the option of resting or watching silent, sub-titled video. They could take a break at any time by pressing a button to pause the experiment.

EEG were recorded using the BrainVision ActiCHAMP amplifier and two EP-Preamp ABR preamplifiers. Two bipolar channels were recorded: FCz referenced to each of the earlobes. The ground electrode was placed on the forehead at Fpz. The two channels were averaged during preprocessing to increase SNR and because stimuli were presented diotically.

Triggers for synchronizing stimulus presentation to EEG data were generated using the sound card’s optical digital audio output, converted to TTL voltages using a custom-made trigger box (Maddox, 2020), and then fed to the EEG amplifier. The 0.9 ms earphone tube delay was corrected in preprocessing. Triggers were also used to precisely mark the end of each trial.

### Preprocessing and ABR TRF calculation

Preprocessing and analysis were done in python scripts that made heavy use of the mne package (Gramfort et al., 2013; mne Developers, 2024). Raw data were high-and low-pass filtered at 1 and 2000 Hz using first order causal infinite impulse response filters (IIR), and then notch filtered at odd multiples of 60 Hz, also with causal IIR filters, to remove power line noise. EEG epochs were created for each trial that began 1 s before stimulus start and ended 1 s after stimulus end. ABR TRFs were calculated from each epoch through frequency-domain deconvolution, using the glottal pulse trains as the stimulus regressor, exactly as in Polonenko and Maddox (2021a). ABRs for each participant were calculated for each talker in each epoch and then averaged across talkers and epochs to yield a single response for each SNR condition (see supplementary data for separate responses to each talker). To improve signal quality, each trial’s contribution to the average was weighted by the inverse of the EEG variance during that period, a technique we have shown to be effective in several past studies (Polonenko and Maddox, 2019, 2021b, 2021a). After ABRs were calculated, they were bandpass filtered between 150 and 2000 Hz (causal, first-order, IIR) for better visualization of early ABR waves.

For experiments with longer stimuli, drift between the sound card and EEG clocks can cause smearing of responses. We eliminated this problem by using the triggers that marked the start and end of each trial to calculate the drift and compensate by resampling the glottal pulse train regressor before deconvolution.

### ABR analysis

ABR wave V peaks and troughs were picked automatically by taking the maximum and minimum point of the waveform between 6.8 and 11.5 ms. Before picking, each waveform was low-pass filtered at 500 Hz to reduce noise using a zero-phase filter so that peaks did not shift. All picks were inspected by both authors to confirm validity. Individual participant waveforms for each SNR with labeled peaks can be seen in supplementary data. The wave V amplitude was taken as the difference between the peak and trough voltages. The wave V latency was taken as the time of the peak voltage.

We assessed waveform quality by computing the waveform SNR, not to be confused with stimulus SNR. The signal+noise variance, 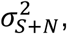 was computed from the response between 4 and 12 ms (a segment duration of 8 ms). The noise variance, 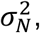 was computed by dividing the pre-stimulus period from the same waveform (which contained only noise), into adjacent 8 ms segments, computing the variance of each, and averaging. The SNR in decibels could then be calculated as

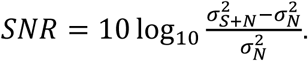

Wave V amplitude and latency changes with stimulus SNR were evaluated with a linear mixed effects model with fixed effects of the logarithm of the number of talkers in the mixture to transform the data for linearity, and random effects of intercept and slope per participant. The random effects for each participant were extracted to determine each participant’s change in the two wave V metrics with stimulus SNR. These slope fits, along with wave V latency and amplitude were tested for a relationship with the behavioral CRM thresholds for peaky speech using Pearson’s *r*.

### Determining minimum necessary experimental conditions

The slopes of the change in wave V amplitude and latency as SNR is decreased represent interesting parameters for future studies of individual differences. To expedite such studies, we determined whether those slopes could be accurately determined from only the extreme SNR conditions, as opposed to all four tested here. We first fit a slope-intercept model to the full dataset for each participant (four points). Next, we did the same for only the Clean and −6 dB conditions (two points). We then computed the squared correlation (variance explained) between the slopes from the full and reduced datasets across participants, with higher numbers indicating a better match (and thus lesser importance of the inner two SNRs). We note that since the number of points in the reduced set for each participant matches the number of parameters in the model (slope and intercept), the resultant model is equivalent to simply “connecting the dots.” We carried out this process for both wave V amplitude and latency.

### Software Accessibility

The software used to analyze the data and generate the figures that appear in this paper can be accessed at https://github.com/polonenkolab/peaky_snr. All experimental data is uploaded to OpenNEURO and can be accessed at https://openneuro.org/datasets/ds005408/versions/1.0.0.

## Results

### Speech-in-noise thresholds are the same for unaltered and peaky speech

Before assessing the neurophysiological effects of speech-on-speech masking, it was important to confirm that the perceptual masking effects for peaky speech matched that of the unaltered speech stimuli. Figure 2 shows the thresholds, expressed in decibels target-to-masker ratio, for each participant as well as the average. The mean ± SEM thresholds were 1.20 ± 0.31 and 1.38 ± 0.33 dB for unaltered and peaky speech, respectively. This difference of 0.18 ± 0.02 dB was not significant (paired t-test; *t*(*24*) = −0.92, *p* = 0.37).

**Figure 1.**
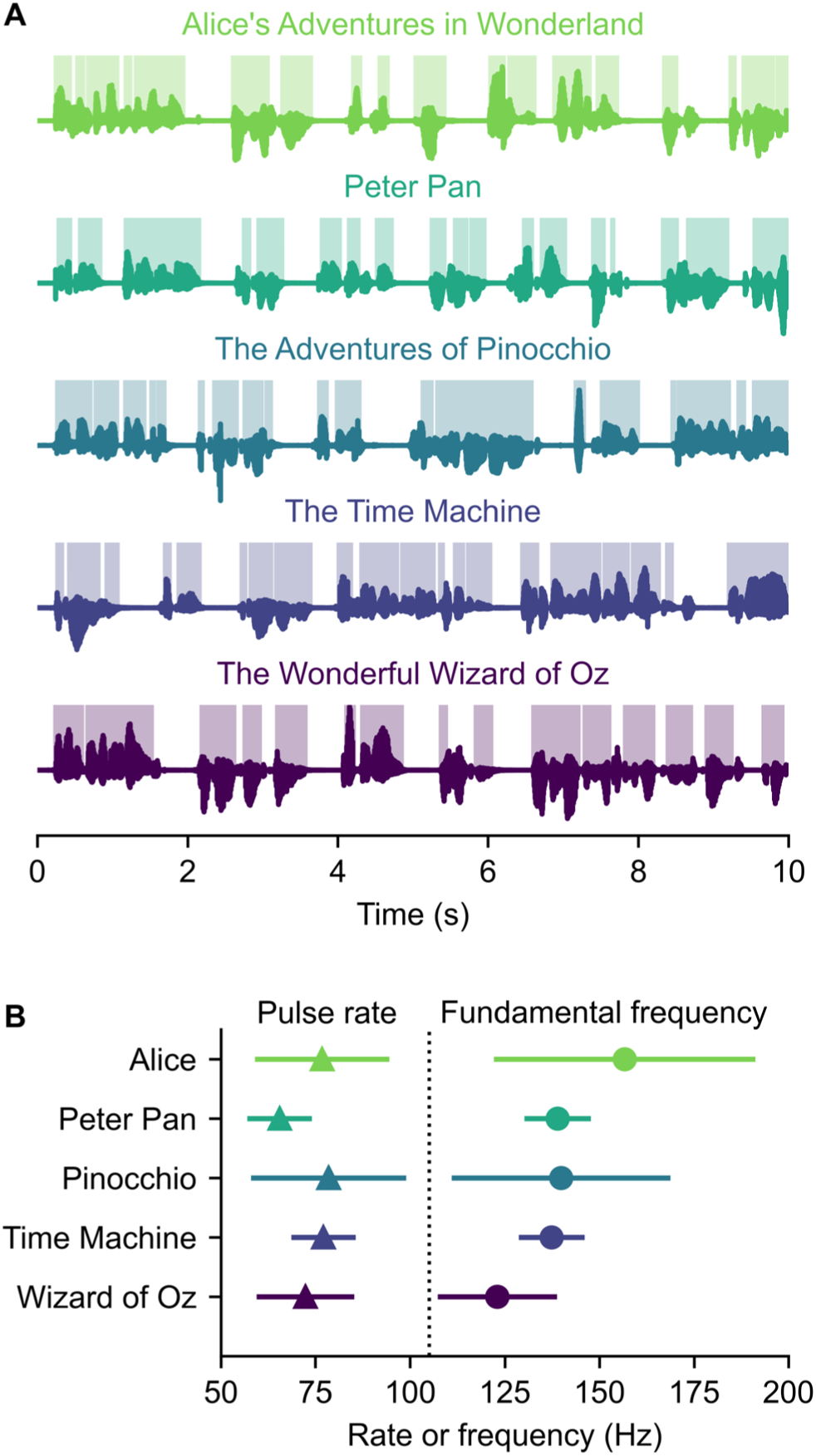
Experimental stimuli. A) An example 10 seconds from each of the five audiobooks. Dark lines show waveforms, with paler highlighted regions corresponding to voiced portions of speech (i.e., where there were glottal pulses). B) The overall glottal pulse rate (number of pulses per second) and average fundamental frequency (number of pulses per second of voiced speech) of each audiobook.

**Figure 2.**
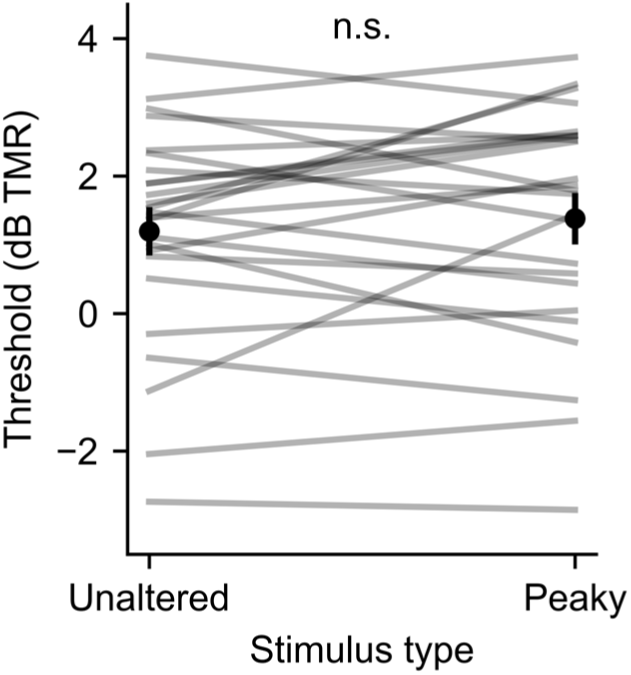
Comparison of speech reception thresholds for unaltered speech versus resynthesized peaky speech. Gray lines show individual participants. Black points and error bars show mean and ±1 SEM.

### ABRs change systematically with worsening SNR

Clear ABRs were present for all SNR conditions (Figure 3). Even though early ABR waves I and III were present in most participants (visible at ∼3.5 and 5.5 ms in Figure 3), they were less distinct than wave V, which makes identifying trends in small differences across SNR conditions difficult. As such, we focused on SNR-dependent changes to the amplitude and latency of wave V.

**Figure 3.**
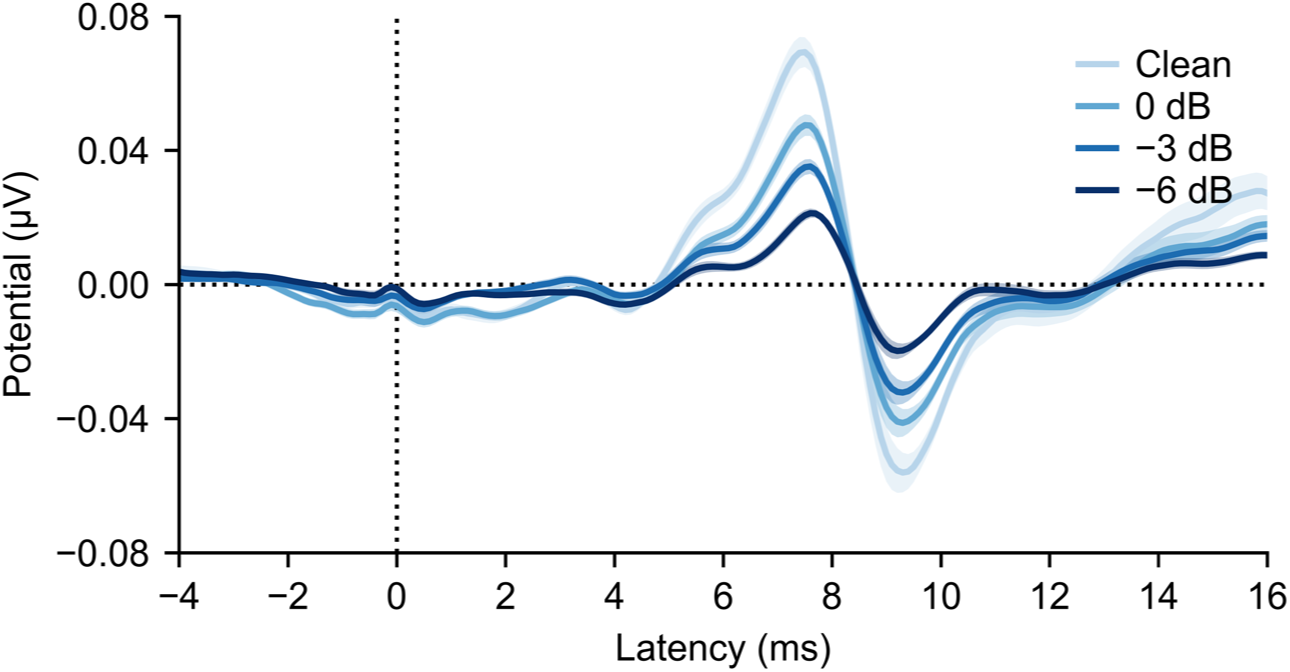
Grand average waveforms across 25 participants for each SNR (darker colors indicate lower SNR). Wave V is the prominent peak at ∼7.5 ms. Shaded areas show ±1 SEM around the mean.

There was a pronounced and consistent reduction of wave V amplitude with decreasing SNR for every participant (Figure 4A). The mean amplitude for the clean speech was 0.15 μV, shrinking to 0.10, 0.08, and 0.05 μV for the 0, −3, and −6 dB SNR conditions, respectively. An ANOVA following a linear mixed-effects model of wave V amplitude with SNR as a fixed effect and participant as a random effect shows a strong, significant effect of SNR (*F*(1, 24) = 337, *p* = 1×10^−15^, *η_p_^2^* = 0.93; estimated intercept ± SEM: 0.146 ± 0.007 µV, estimated slope ± SEM:-0.041 ± 0.002 µV / log_2_(# talkers)).

**Figure 4.**
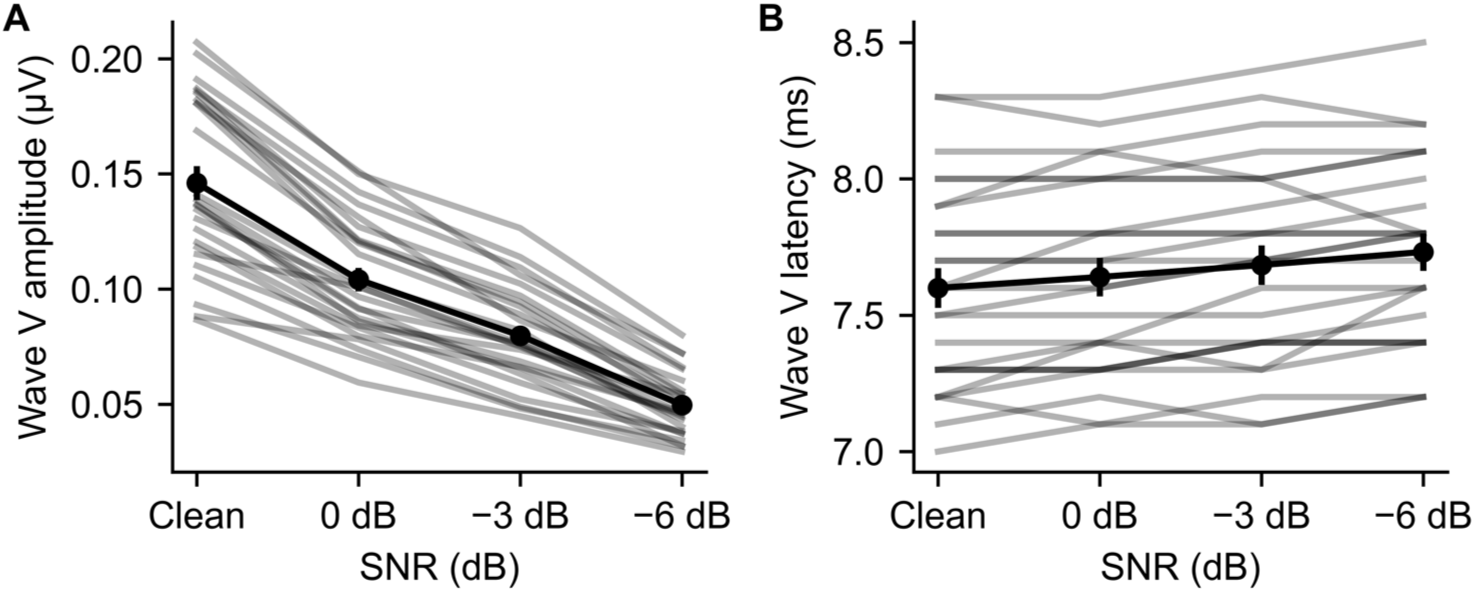
Wave V amplitude (A) and latency (B) across SNR conditions. Gray lines show individual participants. Black points and error bars show mean and ±1 SEM.

Wave V latency increased with worsening SNR (Figure 4B). The mean latency for clean speech was 7.60 ms, increasing to 7.64, 7.68, and 7.73 ms for 0, −3, and −6 dB SNR. Even though there was more variation across SNR in the individual participant latencies than for amplitudes, all 25 participants showed an overall increase in latency when fit with a line. An ANOVA with the same structure as above shows a significant effect of SNR on wave V latency (*F*(1, 24) = 39.664, *p* = 2×10^−6^, *η_p_^2^* = 0.62; estimated intercept ± SEM: 7.593 ± 0.075 ms, estimated slope ± SEM: 0.057 ± 0.009 ms / log_2_(# talkers)).

### Responses are quick to record across stimulus SNR conditions

Recording time is a limiting factor in every electrophysiological experiment, so we consider that next. A recording can be considered complete when the quality of the response reaches a certain threshold—in this case, we calculate the point in the acquisition at which the response waveform reaches an SNR of 0 dB, consistent with many prior studies (note that this response SNR is distinct from the speech SNR that comprises this experiment’s primary independent variable). Recording time cumulative distributions are shown in Figure 5A for each speech SNR condition. For clean, 0 dB, and −3 dB conditions, 90% of recordings took approximately 3 minutes, and all were under 5 minutes. For the −6 dB condition, 90% and 100% of recordings were completed in 8 and 9 minutes, respectively.

**Figure 5.**
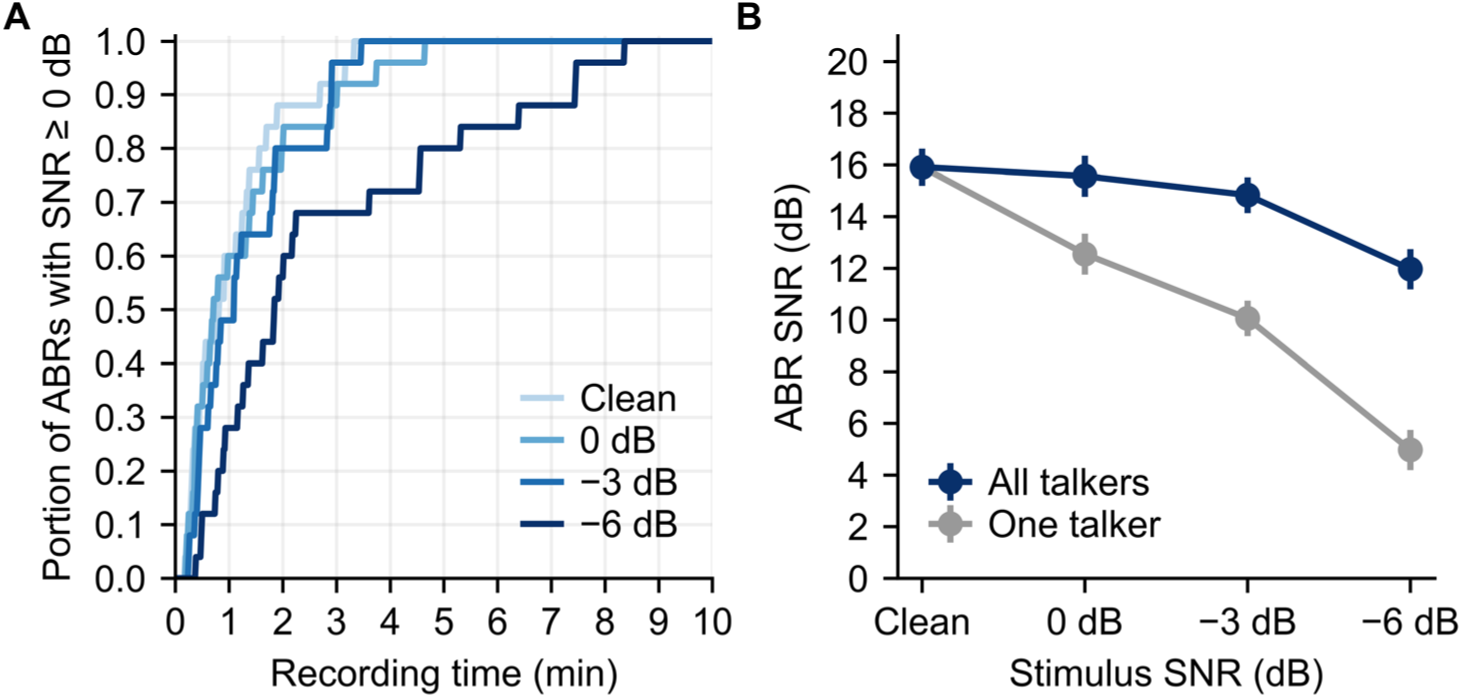
A) Cumulative distributions of recording time for each participant’s ABR to reach at least 0 dB SNR for each stimulus SNR condition. B) Grand average ABR waveform SNR (computed in the 4–12 ms region) for each stimulus SNR condition using all recorded data (30 minutes per condition). Dark line indicates using all talkers and averaging (e.g., five talkers in the −6 dB condition), with light gray indicating what the SNR would be if only one talker were considered the target. Error bars show ±1 SEM.

The waveform SNR (which is the primary determinant of recording time) is shown in dark line in Figure 5B. The average SNRs were 15.8, 15.5, 14.7, and 11.8 for the clean through −6 dB conditions after the entire recording time of 30 minutes each.

The SNR difference of 4 dB between the two extreme conditions (Clean and −6 dB) is smaller than might be expected *prima facie* given the substantial two-thirds reduction in wave V amplitude between them. The reason for this discrepancy is that, even though the actual recording time is 30 minutes for both conditions, the duration of speech stimuli presented and used to calculate the responses was not. In the Clean condition (one talker), there were 30 minutes of speech presented in the 30-minute recording. For the −6 dB condition, there were five concurrent talkers, meaning that 150 minutes of speech stimulus were presented in the 30-minute recording. A fivefold increase in data leads to a noise reduction (and SNR improvement) of 7 dB. The 0 and −3 dB stimulus conditions benefit similarly from the recording scheme, offering waveform noise reductions of 3 and 5 dB. The light gray line in Figure 5B shows what the SNR would be if only one masked talker was presented at a time.

We used four SNR conditions here so that smooth changes in ABR morphologies could be observed, but recording time could be further reduced by running a subset of conditions. To determine such a scheme’s viability, we determined how well the change in wave V amplitude and latency across SNR could be predicted from only the extreme SNRs (Clean and −6 dB) instead of all four datapoints for each participant (see Methods for details). For wave V amplitude, the variance explained between the full and reduced datasets was 99.7%. For latency it was 97%. Both of these are strikingly high, indicating that the intermediate SNRs offer essentially no additional information for computing the slopes that isn’t already present in the extreme conditions. When we follow the same procedure but keep only the inner SNRs instead of the extremes, the variance explained by the reduced model is only 36% and 8% for wave V amplitude and latency, respectively, indicating that the prior numbers are not artificially high as a result of including common datapoints in both models. Thus, for experiments (or diagnostics) where the variation in wave V amplitude and latency across stimulus SNR is of interest, two conditions may provide essentially the same information that four do.

### Exploratory analyses find no correlation between ABR and behavioral speech-in-noise thresholds

The relationship between ABR changes with stimulus SNR and perception are of great interest for future work, but were not a focus of this study. Still, because we tested speech reception thresholds before the ABR experiment, we performed an exploratory analysis to see if they were related. The results are plotted in Figure 6. The correlation between wave V amplitude change and speech reception threshold was not significant (*r* = 0.04, *p* = 0.85). The correlation between wave V latency change and speech reception threshold was also not significant (*r* = 0.33, *p* = 0.11). It is important to note that the participants tested in this experiment were young listeners with normal hearing thresholds, and we further have no reason to believe they struggle with listening in noise. There is thus very little variance to explain. Future studies will require recruitment of participants that span the range of whatever aspect of auditory function is of interest.

**Figure 6.**
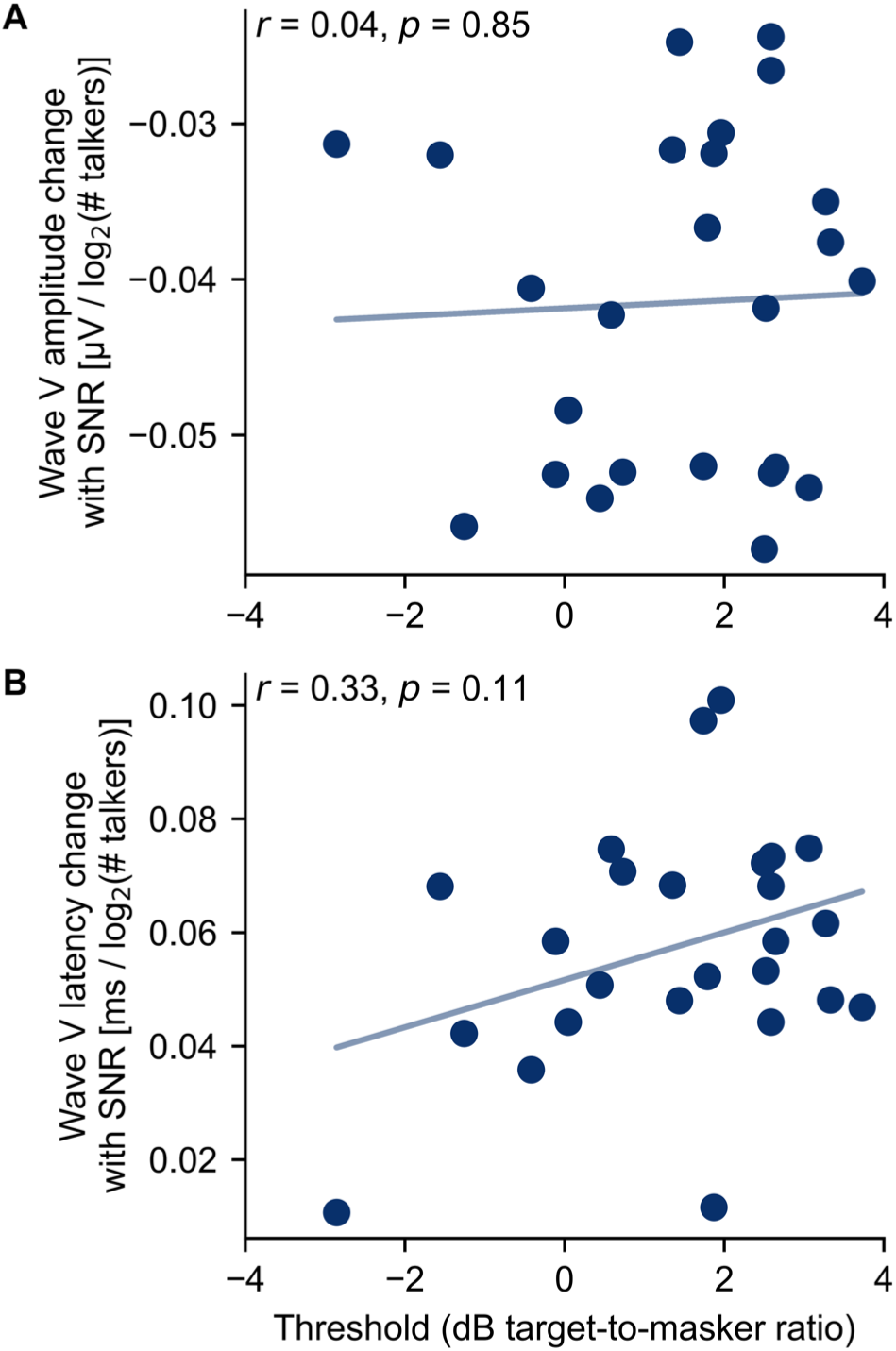
The relationship of speech-in-noise thresholds with the slope of the wave V amplitude change (A) and latency change (B) across SNR. Individual participants are plotted as points along with the best linear fit. There was no significant correlation between either wave V parameter and speech-in-noise perception.

## Discussion

This study aimed to understand how the subcortical response to continuous speech changes under energetic masking by other speech. This is an important question to address because understanding speech in noisy scenarios is both a common and frustrating experience for many people. While behavioral studies of masking are plentiful, physiological studies aimed at the subcortex are far fewer in number and have faced limitations. Several studies have measured the frequency-following response to a repeated syllable presented in babble noise (e.g., Wong et al., 2007). Those responses, however, are generated by an unknown mixture of sources, including both subcortical and cortical areas (Coffey et al., 2016, 2017, 2019). Other studies have measured a canonical ABR with clear generators, but have used artificial stimuli such as clicks that are far removed from natural speech (e.g., Mehraei et al., 2016). In this study, we utilized the peaky speech TRF paradigm (Polonenko and Maddox, 2021a), which allows clear ABRs to be measured from naturalistic speech stimuli (here, audiobooks).

### Effects of speech masking on subcortical continuous speech encoding

We observed two significant effects of SNR on the encoding of individual speech streams: the wave V component of the ABR decreases in amplitude and increases in latency as the number of competing streams increases. These changes were quantified by extracting amplitude and latency parameters of individual response waveforms (Figure 4, with individual responses shown in supplementary data) as well as the grand averaged waveforms (Figure 3). These changes in wave V are similar to a recent study that examined the effect of sound level of a single speech stream using analysis methods much like ours (Kulasingham et al., 2024b). Thus, two distinct factors that can make speech more difficult to understand (either reducing its energy or increasing the energy of competing sounds, thereby decreasing the SNR) both exert similar effects on the subcortical response waveform.

The focus on ABR wave V here (and in other papers) is mostly one of convenience: wave V is a large and distinct peak in each participant’s response. The ABR waveform has additional components originating from the ascending pathway’s earlier neurogenerators (wave I: the small bump at ∼3 ms and wave III: the bump at ∼6 ms on the rise to the wave V peak) that can be seen in the grand average waveforms. Wave III, which is generated by the cochlear nucleus (Burkard et al., 2006 p.5), seems to show a similar amplitude change as wave V without much change in latency, but the component was not robust enough for trends to be analyzed at the level of single-participant waveforms. Wave I, generated by the auditory nerve, was similarly too small for detailed analysis, even though it was present in the grand averages. The compound action potential (CAP) is a response with the same generator as wave I that is measured with an electrode placed on the eardrum (Simpson et al., 2020). It is substantially larger than wave I with much better SNR, but the placement of the electrode adds complication to running the experiment and was not pursued here.

There are many possible stimulus regressors for computing the subcortical TRF with different strengths and weaknesses (Kulasingham et al., 2024a). Recent work from our lab directly compared the peaky speech paradigm’s glottal pulse regressor to a different regressor based on the predicted firing rate of an auditory nerve model and found that the latter yields slightly higher SNR (Shan and Maddox, 2024). We opted to use the glottal pulses here because, unlike the auditory nerve model, it provides amplitudes in physical units (µV) and the latencies of the responses do not require adjustment to compensate for the estimated model delay (Shan et al., 2024). Responses computed with the auditory nerve model regressor are included as supplementary data.

### Practicality for clinical measurement

In addition to investigating the overall effects of SNR on brainstem responses, we are also interested in adapting this paradigm for potential clinical use. Neuropathy and synaptopathy of the auditory nerve have been hypothesized to play a role in poorer speech-in-noise perception in people with normal audiologic pure-tone thresholds (Bharadwaj et al., 2014). Finding an indicator of disordered neural processing that is reliable across individuals would be highly valuable, though has proven difficult so far (Prendergast et al., 2015; Plack et al., 2016). Wave I (which was present in our data but too small to show trends) is often the response of interest, since it indexes auditory nerve activity (Grant et al., 2020). To evoke robust wave I responses, measurements are done with very high-level clicks, but using an eardrum electrode to measure the auditory nerve CAP allows lower-level stimuli to be used (Simpson et al., 2020). An ecologically relevant paradigm that measures the CAP evoked by a speech mixture at a conversational listening level may provide a more informative clinical measure.

In addition to the effects that masking has on subcortical speech responses, we also assessed how quickly those effects could be estimated. Responses for the clean, 0, and −3 dB conditions reached the criterion SNR of 0 dB in under 5 minutes for all participants, with the −6 dB condition reaching criterion in under 10 minutes for all participants (Figure 5). The median time to reach criterion was under 2 minutes for all SNR conditions. Since the wave V amplitude and latency showed consistent changes as SNR decreased, a clinical test based on the slope of those changes could be sped up by reducing the number of responses measured. We found that using only the Clean and −6 dB conditions to compute the slopes provided essentially all the information present in the complete dataset. With only two conditions needed, a simple difference replaces computing the slope, leading to metrics that are quick to measure and simple to calculate.

We also performed a behavioral study of speech recognition thresholds in noise. Its primary purpose was to confirm peaky speech showed the same intelligibility as unaltered natural speech (which it did, Figure 2), but it also allowed an initial exploratory analysis of whether the SNR-driven changes to the ABR were correlated with behavioral thresholds. That analysis showed no relationship between wave V amplitude or latency changes with the behavioral results. We caution, however, that this was really an “analysis of convenience.” Participants were all young with normal hearing and were not recruited based on reported listening difficulties or a history of noise exposure, so there was very little variance to explain in the first place. Future studies could focus on people with hearing loss or people with normal thresholds who report listening challenges. Even if in those populations, too, no correlations were observed across individual participants, determining the overall effects of SNR on subcortical speech coding in those groups would be novel and interesting to compare to the present paper’s findings.

### Why cortical responses were not studied here

TRFs are frequently used to assess sound encoding in the cortex, and can be informative about responses driven by basic acoustics or more complicated aspects of an input stimulus such as phonetic features (Di Liberto et al., 2023) or semantic surprisal (Broderick et al., 2018). The focus of the current investigation was the effect of masking on subcortical encoding, but we have previously shown that the same TRF methods used here will also provide cortical responses, simply by extending the end of the analysis window later and using different offline filter parameters. We did not analyze them here because several uncontrolled experimental factors could have affected cortical responses. First, participant arousal state was not controlled. Some participants were alert while others may have dozed off. Sleep has a suppressive effect on cortical evoked potentials (Deiber et al., 1989), but does not appreciably affect subcortical responses (Campbell and Bartoli, 1986)—in fact, a sleeping patient is ideal when subcortical responses are used in a diagnostic ABR exam.

Second, even though participants were given the option of reading or watching a subtitled movie, we cannot be certain that they didn’t decide to listen to excerpts of specific stories (or focus on specific talkers) when they were part of a presented stimulus. Even though that was unlikely given the stimulus randomization, directed attention strongly affects cortical responses (O’Sullivan et al., 2014), while its effect on subcortical responses is contested (Varghese et al., 2015) and generally small, even in studies that report it. Interestingly, prior work has shown that the encoding of both the attended and unattended speech streams are relatively stable (with the attended stimulus showing stronger responses) across a wide range of target-to-masker level ratios (Ding and Simon, 2012). Such a study demonstrates the importance of explicitly controlling for attention when investigating cortical encoding of multiple sound sources.

## Acknowledgements

Yathida Melody Anankul assisted with participant recruitment and data collection. This work was supported by two grants: NSF CAREER 2142612 and NIH R01DC017962.

## Supplementary Data

**Figure S1.**
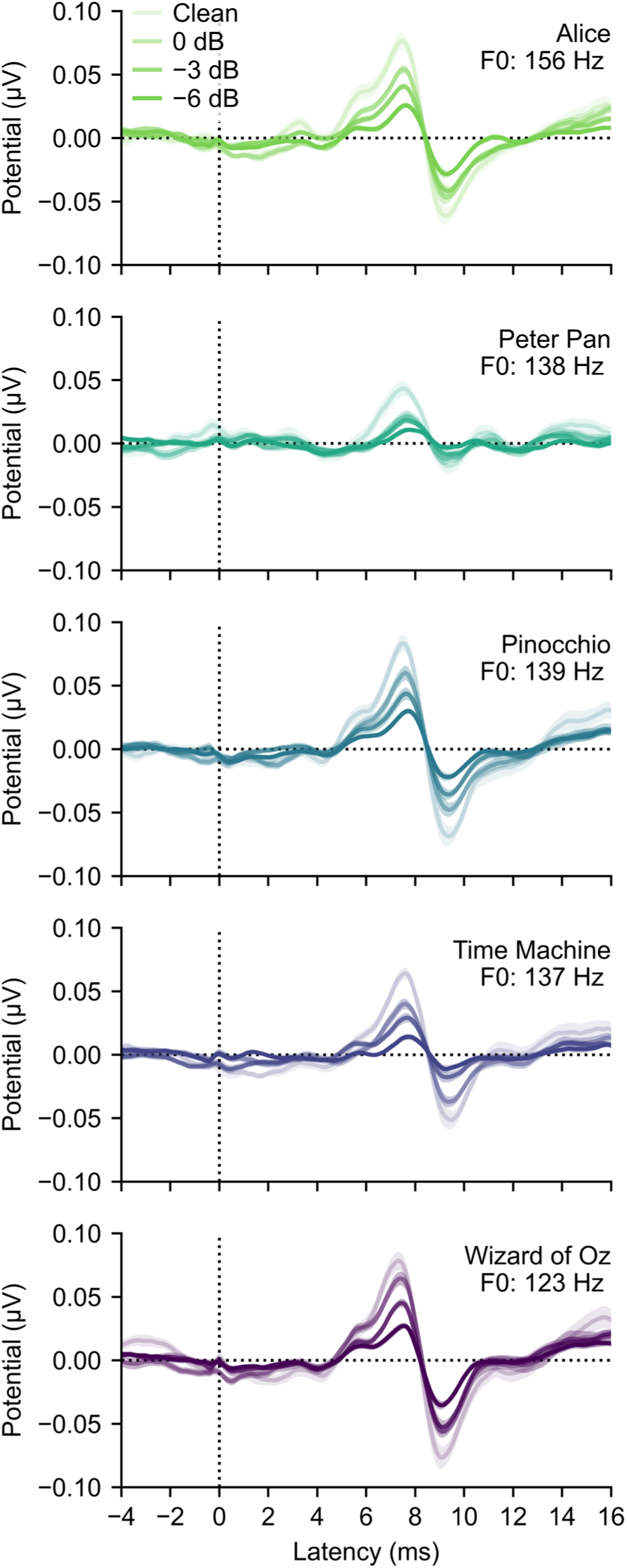
Grand average waveforms in response to each of the five stories. Plotted as in Figure 3.

**Figure S2.**
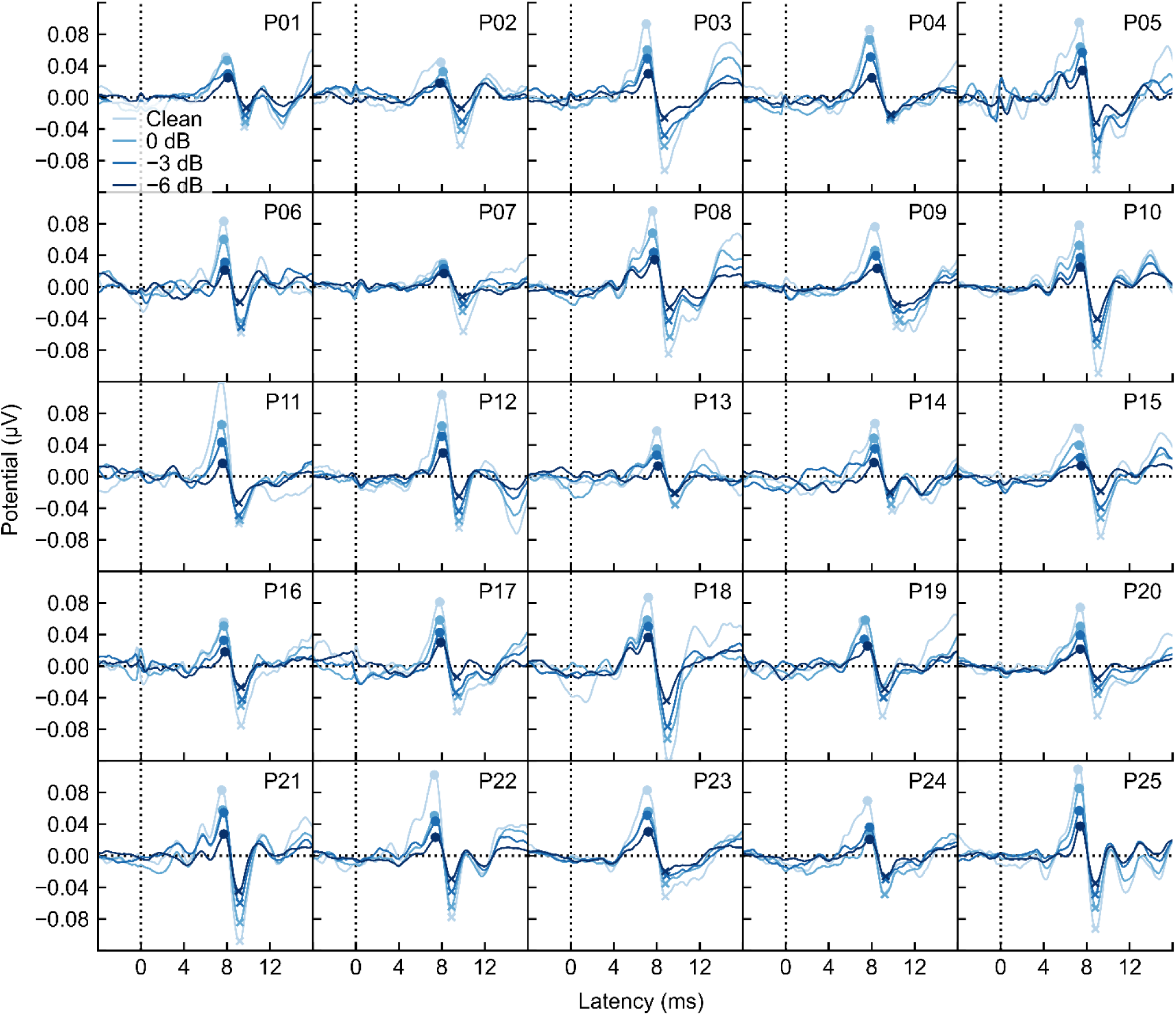
Individual participants’ responses for each SNR condition. Wave V peaks and troughs are marked with dots and exes, respectively.

**Figure S3.**
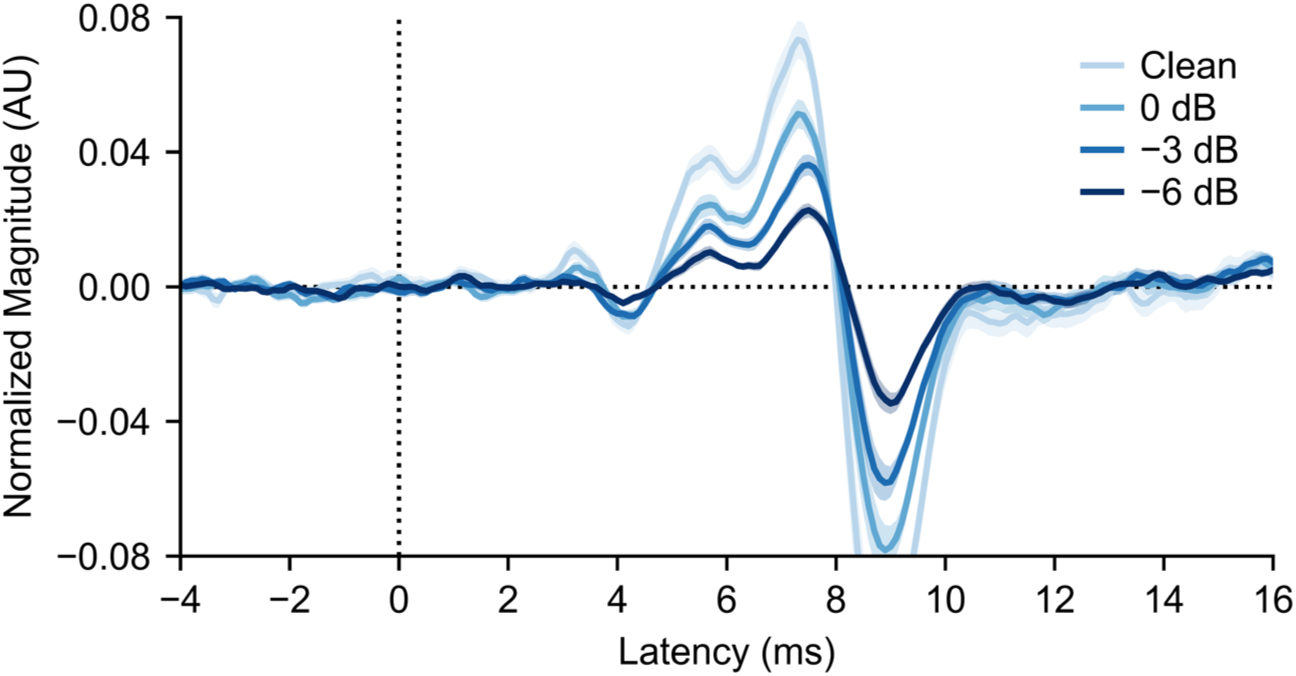
Grand average waveforms generated from the auditory nerve model regressor (Shan et al., 2024). Plotted as in Figure 3.

**Figure S4.**
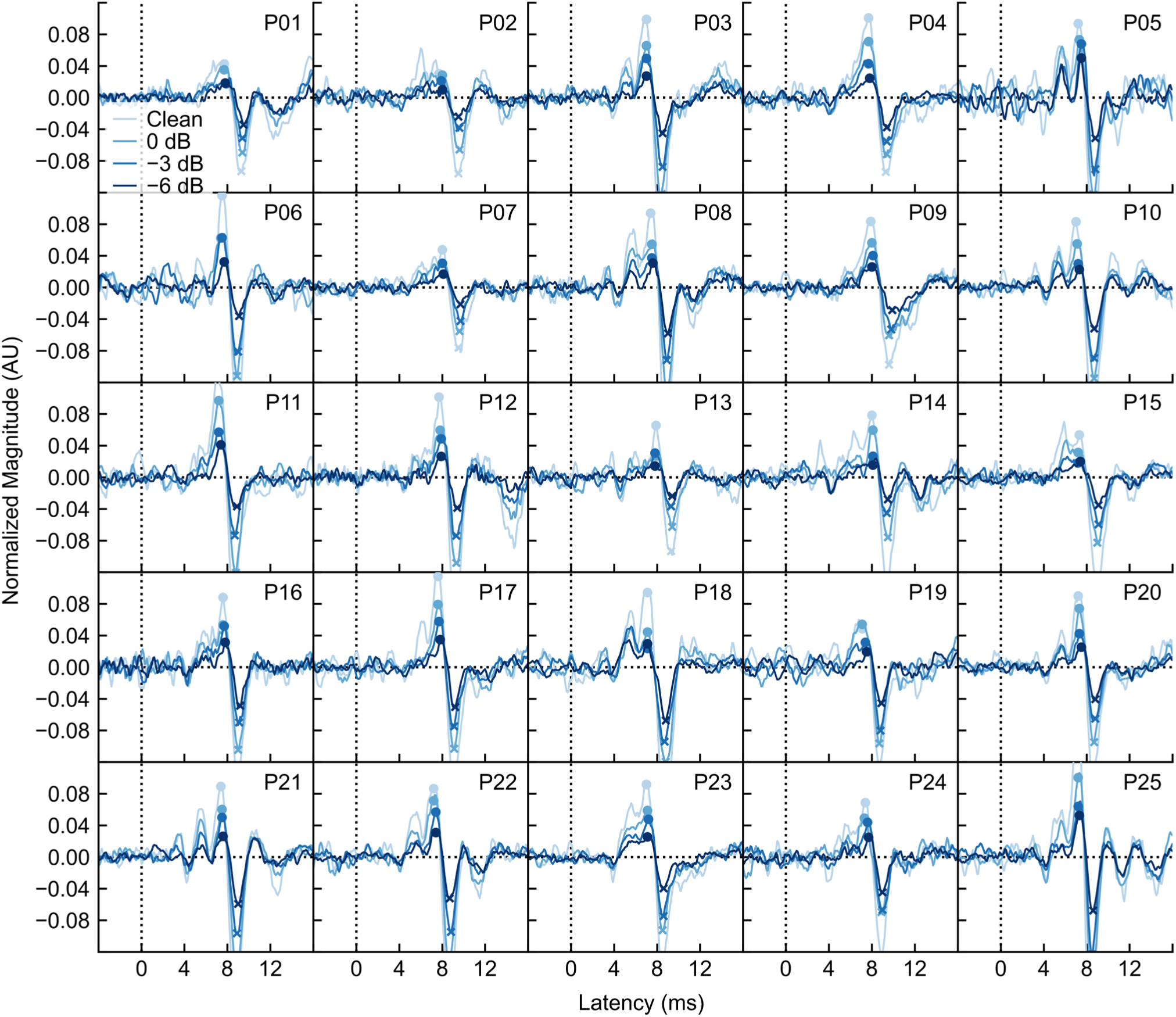
Individual participants’ responses for each SNR condition, using the auditory nerve model regressor (Shan et al., 2024). Wave V peaks and troughs are marked with dots and exes, respectively.

